# Elastin levels are higher in healing tendons than in intact tendons and influence tissue compliance

**DOI:** 10.1101/2020.05.26.065433

**Authors:** Anna Svärd, Malin Hammerman, Pernilla Eliasson

## Abstract

Elastic fibers containing elastin play an important role in tendon functionality, but the knowledge on presence and function of elastin during tendon healing is limited. The aim of this study was to investigate elastin content and distribution in intact and healing Achilles tendons and to understand how elastin influence the viscoelastic properties of tendons. The right Achilles tendon was completely transected in 81 Sprague-Dawley rats. Elastin content was quantified in intact and healing tendons (7, 14, and 28 days post-surgery) and elastin distribution was visualized by immunohistochemistry. Degradation of elastin by elastase incubation was used to study the role of elastin on viscoelastic properties. Mechanical testing was either performed as a cyclic test (20x 10 N) or as a creep test. We found significantly higher levels of elastin in healing tendons at all time-points compared to intact tendons (4% in healing tendons 28 days post-surgery vs 2% in intact tendons). The elastin was more widely distributed throughout the extracellular matrix in the healing tendons in contrast to the intact tendon where the distribution was not so pronounced. Elastase incubation reduced the elastin levels by approximately 30% and led to a 40-50% reduction in creep. This reduction was seen in both intact and healing tendons. Our results show that healing tendons contain more elastin and is more compliable than intact tendons. The role of elastin in tendon healing and tissue compliance indicate a protective role of elastic fibers to prevent re-injuries during early tendon healing.

**PLAIN LANGUAGE SUMMARY:** Tendons transfer high loads from muscles to bones during locomotion. They are primarily made by the protein collagen, a protein that provide strength to the tissues. Besides collagen, tendons also contain other building blocks such as e.g. elastic fibers. Elastic fibers contain elastin and elastin is important for the extensibility of the tendon. When a tendon is injured and ruptured the tissue heals through scar formation. This scar tissue is different from a normal intact tendon and it is important to understand how the tendons heal. Little is known about the presence and function of elastin during healing of tendon injuries. We have shown, in animal experiments, that healing tendons have higher amounts of elastin compared to intact tendons. The elastin is also spread throughout the tissue. When we reduced the levels of this protein, we discovered altered mechanical properties of the tendon. The healing tendon can normally extend quite a lot, but after elastin removal this extensibility was less obvious. The ability of the healing tissue to extend is probably important to protect the tendon from re-injuries during the first months after rupture. We therefore propose that the tendons heal with a large amount of elastin to prevent re-ruptures during early locomotion.

## INTRODUCTION

Elastic fibers are composed by the extracellular protein elastin and microfibrils consisting of fibrillin-1 coupled with microfibril associated protein 4 (1). Elastin is an insoluble polymer made from the monomeric soluble precursor tropoelastin (2). The tropoelastin monomers are linked by amino acids desmosines (3, 4) through action of enzyme lysyl oxidase (5–7). Fibrillin-1 surrounds the elastin and has been suggested to guide elastogenesis by providing a scaffold for elastin. Other proteins like fibulin-5 have also been shown to be important for cross-linking the elastic fiber network (8). Elastic fibers can be stretched and recoil numerous times and thereby play an important role in the function of e.g. arterial walls, lungs, skin, and tendons (9, 10). In tendons, elastin constitute of 1-5% of the dry weight and is found throughout the tissue, especially between collagen fascicles in the interfascicular matrix and in the pericellular environment (9, 11, 12).

The role of elastin in tendons and ligaments have previously been studied by using elastase digestion treatment. The elastase cleaves the tropoelastin and destroys the elastin network but leaves the crosslinks intact (13). By destroying the network with elastase, the function of the elastic fibers is impaired (14). Studies using elastase have shown that elastin seems to be important for several mechanical properties, especially in response to shear forces and at low tensile forces. By degrading elastin in medial collateral ligaments, stress up to 2 MPa was decreased while peak strength, peak strain, modulus and hysteresis were unaffected (14, 15). However, rat tail tendon fascicles had a decreased tensile failure strength and strain after elastin degradation, but again no effect on modulus (16). Albeit there are studies on the role of elastin on tendon mechanical properties, these studies are performed on intact tendons. Hence, the knowledge on presence and function of elastin during tendon healing is limited. Furthermore, a deeper knowledge on the role of elastin in specific mechanical properties, such as the viscoelastic properties is also favorable.

The aim of this study was to investigate elastin content and distribution in intact and healing Achilles tendons at various time points after tendon transection and to understand how elastin influence tendon viscoelastic properties. We used a rat model with complete Achilles tendon transection followed by elastin degradation by elastase incubation. We hypothesized that elastin content would be higher in intact tendons compared to healing tendons and that elastin content in healing tendons would increase with time after surgery. We also hypothesized that higher levels of elastin would be linked to more tendon creep as well as more stiffness of the toe-region but not in the linear region of the load-deformation curve.

## MATERIALS & METHODS

### Study design

This study was divided into 3 separate experiments (Table 1). Experiment 1 was performed to investigate levels and distribution of elastin in intact and healing Achilles tendons in relation to mechanical properties. Cyclic mechanical testing on intact and healing tendons (7, 14, and 28 days after tendon transection) was done followed by elastin quantification by a Fastin^®^ assay. Additionally, 2 rats were used for immunohistochemical staining of elastin on intact and healing tendons (14 days post-surgery). Experiment 2 and 3 were performed to study the role of elastin on viscoelastic properties of intact and healing tendons (14 days post-surgery). This was done by enzymatical degradation of elastin by elastase treatment prior to mechanical testing. Cyclic mechanical testing was performed in Experiment 2 and creep test was performed in Experiment 3.

**Table 1:** Experimental setup displaying n-values of tendon samples for treatment and analysis method.

| Exp. | Tendon sample | Treatment |  |  |  | Analysis |  |  |
| --- | --- | --- | --- | --- | --- | --- | --- | --- |
|  |  | Untreated | Control (PBS) | Elastase 2U/ml | Elastase 4U/ml | IHC | Cyclic test | Creep test |
| 1 | Intact | 17 | - | - | - | 2 | 15 | - |
|  | Healing day 7 | 10 | - | - | - | - | 10 | - |
|  | Healing day 14 | 12 | - | - | - | 2 | 10 | - |
|  | Healing day 28 | 10 | - | - | - | - | 10 | - |
| 2 | Intact | - | 9 (14h) | 10 (14h) | - | - | 19 | - |
|  | Healing day 14 | - | 9 (14h) | 10 (14h) | - | - | 19 | - |
| 3 | Intact | - | 10 (14h) | 10 (14h) | - | - | - | 20 |
|  | Healing day 14 | - | 10 (38h) | - | 10 (14h)<br>10 (38h) | - | - | 30 |

- indicates that means were not performed on these tendons.

### Animal handling and surgery

In total, 81 Sprague-Dawley female rats (Janvier, Saint-Berthevin Cedex, France) were used. Thirty-two rats (280 grams SD 15) were used in Experiment 1. Nineteen rats (298 grams SD 22) were used for Experiment 2. Thirty rats (293 grams SD 23) were used for Experiment 3. Ethical approval was obtained from the Regional Ethics Committee for Animal Experiments in Linköping, Sweden (ethical approval identification number 15-15). Rats were monitored once a day and they had access to water and food ad libitum.

The surgery was performed under sedation with isoflurane (Forene, Abbot Scandinavia, Solna, Sweden) and under aseptic conditions. The skin over the right Achilles tendon was shaved and washed with chlorhexidine ethanol. A small skin incision was made lateral to the Achilles tendon and the tendon complex was exposed. The right Achilles tendon was completely transected transversely while the contralateral tendon was kept intact, as control. The adjacent plantaris tendon was removed prior to transection of the Achilles tendon to avoid possible interference in the elastin measurement and mechanical testing. The transected tendons were left unsutured and allowed to heal spontaneously and the skin was closed with two single sutures. Analgesia (0.045 mg/kg buprenorphine, Temgesic; Scering-Plough, Brussels, Belgium) was administrated subcutaneously prior to surgery and thereafter with 12 hours interval until 48 hours had passed. Rats were also given antibiotics (25 mg/kg oxytetracycline, Engemycin; Intervet, Boxmeer, Holland) subcutaneously prior to surgery. The animals were euthanized by CO_2_ asphyxiation followed by confirmation of death by palpation of heart and visual inspection of mucous membranes. The tendons were dissected free from surrounding tissues and harvested together with the calcaneal bone and the distal part of the triceps surae muscle for mechanical testing, elastin quantification and elastase treatment. Tendons for immunohistochemical staining was harvested without the calcaneal bone.

### Elastin degradation with elastase

Immediately after sacrifice, both intact and healing tendons were harvested and fixed in 6-well plates (coated with a silicone layer in the bottom of the wells to allow needle fixation). The tendons were incubated at 37°C with elastase (Sigma Group, Malmö, Sverige, E1250) as follow; 2 U/ml elastase for 14 hours in Experiment 2; 2 U/ml elastase for 14 hours, 4 U/ml elastase for 14 hours or 4 U/ml elastase for 38 hours in Experiment 3 (Table 1). Control tendons were incubated in filtered phosphate buffered saline (PBS) for 14 or 38 hours. All samples were also incubated with 0.1 mg/ml soybean-trypsin inhibitor (Sigma Group, Malmö, Sverige, T6414) to block decomposition of collagen and 1% penicillin/streptomycin as an antibacterial agent. After incubation, tendons were rinsed with filtered PBS and put on ice until mechanical testing was performed later the same day.

### Mechanical testing

Tendon width, depth, total length and gap length were measured with a digital caliper before mechanical testing. Both intact and healing tendons were measured. In experiment 1 and 2 cyclic loading (20 cycles between 1 to 10 N) was applied on tendons in a material testing apparatus (100R, DDL, Eden Prairie, MN) at a constant speed of 0.1 mm/s. Creep was measured between cycle 2 and cycle 20 (Figure 1). Tendon stiffness was measured at two positions of the force-displacement curve in the 20th cycle. The first position was the toe-region (1 to 1.5 N) and the second position was in the linear region (7 to 9 N). Hysteresis was calculated by measuring the area under the curves for the increasing strains minus the area under the curves for the decreasing strains. In experiment 3 tendons were preconditioned by applying 5 cycles of loading between 1 to 5 N at a rate of 0.1 mm/s after which the tendons were rested on ice for approximately 3 hours in wet condition. For all tendons, creep test was performed by applying 8 N at a rate of 1 mm/s followed by holding the force constant for 300 seconds. Creep was calculated as a percentage between the final and the initial deformation. After the creep test was ended, tendon length was measured again, and the tissue was snap frozen in liquid nitrogen and stored at −20°C.

**Figure 1.**
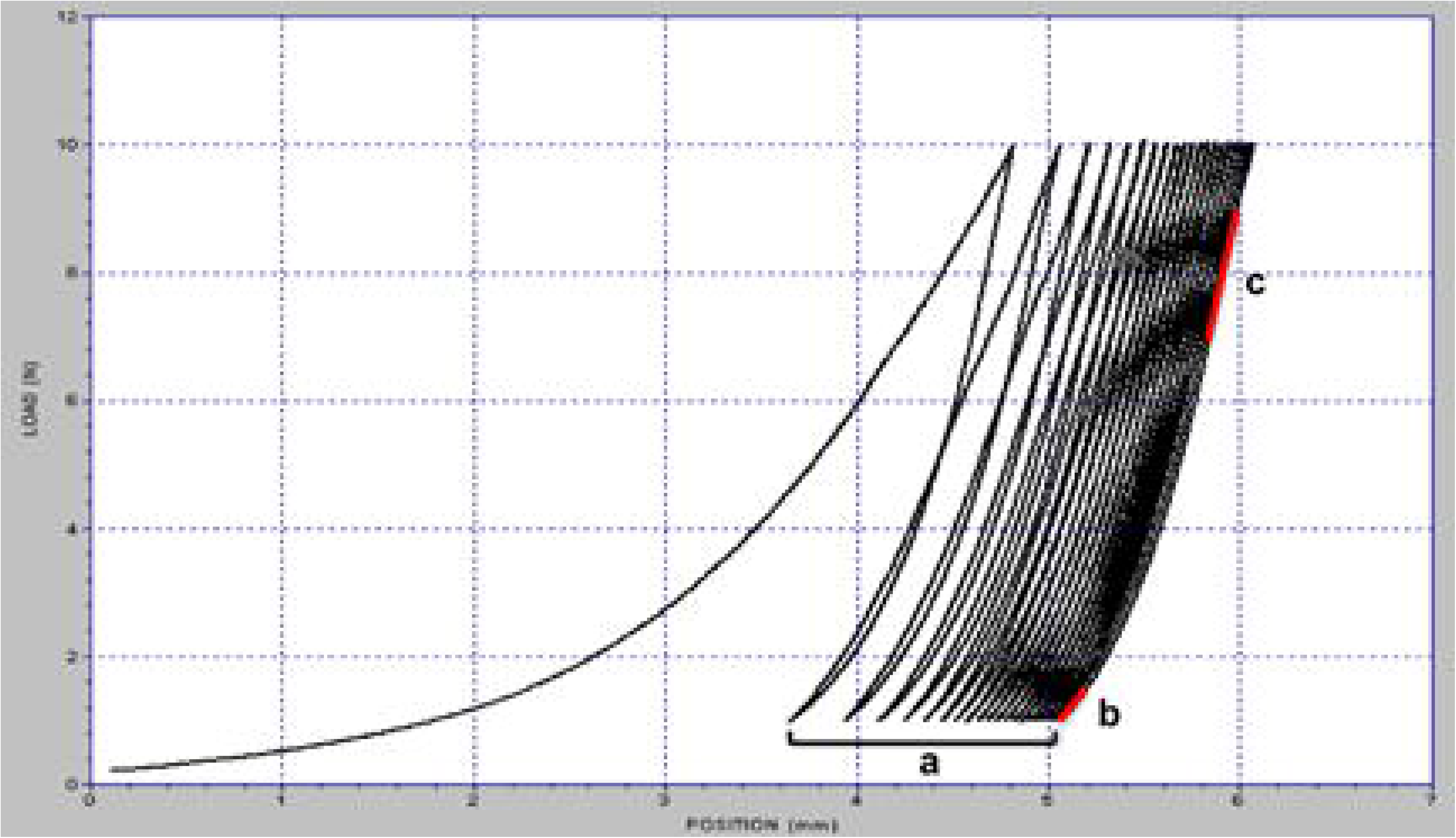
Graph obtained after mechanical testing by cyclic loading (1-10 N, 20 cycles). Creep was measured as difference in length from cycle 2-20 divided by the total length of the tendon (a). Stiffness expressed as Δload/Δposition for toe-region (b) and linear region (c).

In total, 11 tendons were excluded from the analysis of mechanical properties: 4 in Experiment 1, 2 in Experiment 2, and 5 in Experiment 3. The tendons were excluded due to tendon rupture during the test (7 tendons) or due to difficulties with mounting the samples in the mechanical testing machine (3 tendons), or computer error during mechanical testing (1 tendon). These tendons were still used for elastin quantification.

### Elastin quantification

Specimens containing intact tendon tissue or solely newly formed callus tissue (i.e. the tissue between the old tendon stumps) were thawed and trimmed transversely into samples between 10-20 mg. The wet weight was recorded, samples were lyophilized overnight and dry weight was recorded. Measurement of elastin content was performed with a commercial elastin dyebinding assay (Fastin^®^, Bicolor Ltd., Carrickfergus, Country Antrim, UK) according to manufacturer’s instructions. Briefly, samples were digested in 750 μl of oxalic acid (0.25 M) in heating blocks (98°C) for 1 hour. The samples were thereafter cooled down, centrifuged at 10000 rpm for 10 minutes and the supernatants were collected. Additional 750 μl of oxalic acid was added to the residual tissue and the procedure was repeated. All supernatant was sterile filtered, and 250 μl of the supernatant was mixed with a dye, specific for elastin. Samples were agitated for 90 minutes and unbound dye was removed, followed by dissociation of dye-elastin complexes. Samples were then added to a 96-well plate and the absorbance was measured at 513 nm in a plate reader (Epoch 2 Microplate Spectrophotometer, BioTek). A standard curve of known elastin concentrations was used to quantify elastin content in test samples with linear regression. Elastin content was normalized to dry weight of the tendons.

### Immunohistochemical staining

Harvested tendons were kept in 15% sucrose solution (Sigma Group, Malmö, Sverige, S0389), followed by 30% sucrose solution, and finally a mix of OCT (VWR International AB, Spånga, Sweden, 00411243)/30% sucrose solution (1:1). The tendons were snap-frozen in optimal cutting temperature compound (OCT) with liquid nitrogen and stored in −80°C. Frozen tendons were sectioned longitudinally to 7 μm thick slices, comprising the full length of the tendon, and stained for elastin. Sections were hydrated in PBTD (PBS containing 0.1% Tween20 and 1% DMSO) and fixed in 4% formaldehyde for 10 minutes. A blocking buffer containing 5% normal goat serum (Sigma Group, Malmö, Sverige; G9023) diluted in PBTD was added for 2 hours followed by an anti-elastin rabbit polyclonal antibody diluted in PBTD 1:100 (Abcam, Cambridge, UK; ab21610). The samples were incubated with the primary antibody for 18 hours at 4°C followed by 30 minutes washing in PBTD. A goat anti-rabbit IgG secondary antibody (Alexa Fluor Plus 488 Highly Cross-Adsorbed Secondary Antibody; 1:400 diluted in PBTD; Fisher Scientific, Stockholm, Sweden; A32731) was added for 2 hours followed by washing in PBTD for 25 minutes. DAPI (4’,6-Diamidino-2-Phenylindole Dihydrochloride; 1:2000 diluted in PBTD; Thermo Fisher Scientific, Stockholm, Sweden; 62248) was added for 3 minutes followed by washing in PBTD for 25 minutes and mounting with ProLong gold antifade reagent (Thermo Fisher Scientific, Stockholm Sweden; P36930). The stained samples were examined using an Axio Vert. A1 microscope with a Plan Apo 20X and 40X objective (Zeiss, Oberkochen, Germany). The exposure time was held constant for each color channel regarding magnification and tendon samples (intact or healing tendons). All images were adjusted to the negative control staining, where the primary antibody was omitted, to correct for unspecific antibody detection. Skin from the rat was used as a positive control.

### Statistics

Statistical analyzes were performed with GraphPad Prism 8 software. Experiment 1 was analyzed by paired t-tests between intact and healing tendons and one-way ANOVA between healing tendons at different time-points. Simple linear regression analysis was used to analyze correlations between stiffness and stiffness ratio and elastin levels. Intact treated and control tendons in Experiment 2 and 3 were analyzed by unpaired Student’s t-test. Healing tendons in experiment 2 was also analyzed by unpaired Student’s t-test while a one-way ANOVA was used for healing tendons in Experiment 3. Values of p ≤ 0.05 were considered significant.

## RESULTS

### Healing tendons had higher elastin content compared to intact tendons

The elastin content (percentage of dry weight) was twice as high in healing tendons compared to intact tendons at all time-points, 7-28 days post-surgery (p<0.001, Figure 2). Moreover, there was no difference in elastin content between the three time-points post-surgery. Immunohistochemical staining confirmed a wider distribution of elastin in healing tendons, 14 days post-surgery, compared to intact tendons (Figure 3). Most of the elastin was found in the pericellular environment, but it was also found to some extent in other areas of the extracellular matrix. In intact tendons, the elastin was found in the interfascicular matrix and in the pericellular environment between the collagen fibers, leading to a linear appearance of the elastin with black lines (collagen) in-between. However, healing tendons also had a higher number of cells surrounded by elastin, which were distributed evenly throughout the tissue, leading to a wider distribution of elastin in healing compared to intact tendons.

**Figure 2.**
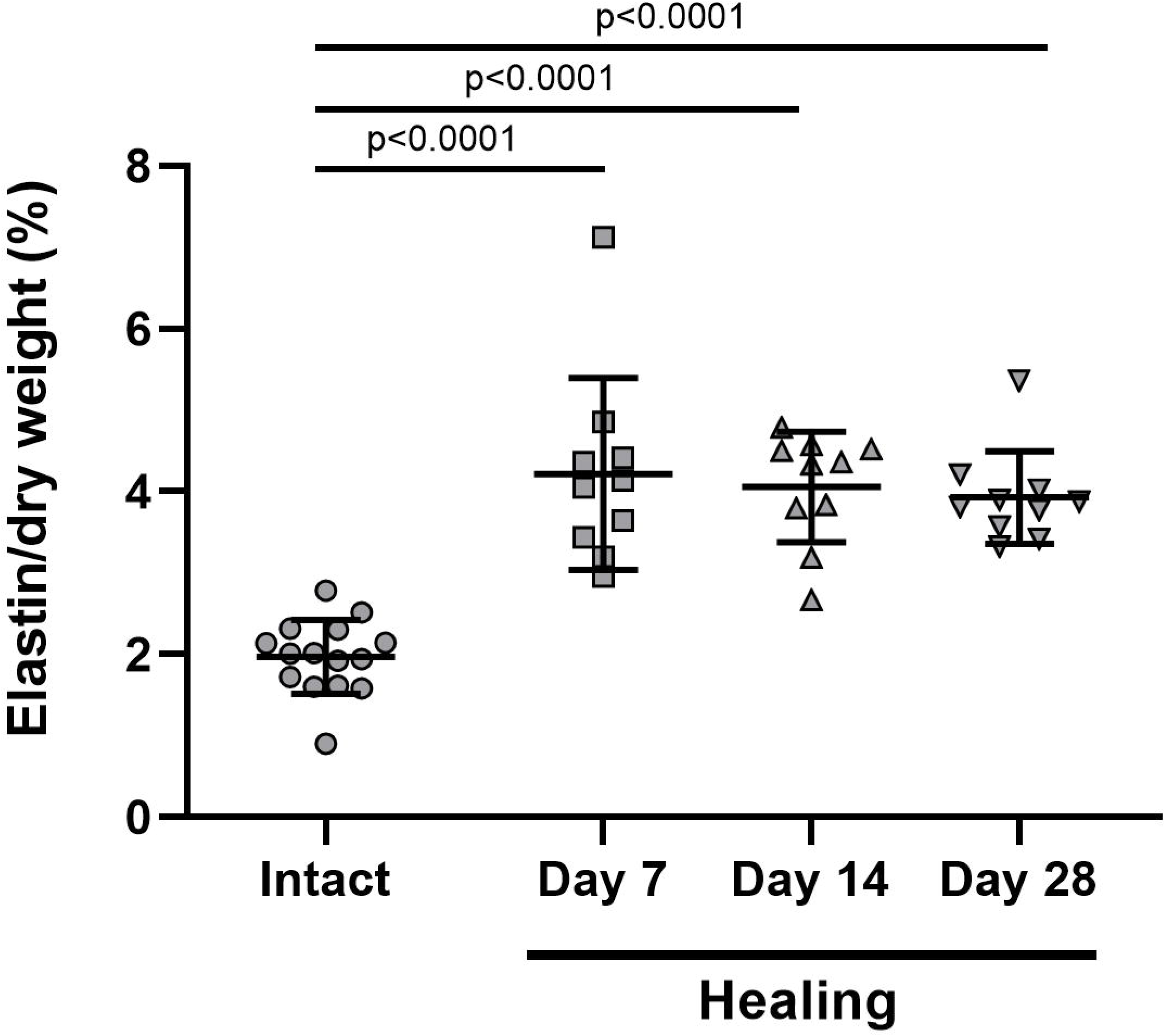
Elastin content in intact and healing tendons. The levels were significantly higher in the healing tendons at 7, 14 and 28 days after tendon transection compared to intact tendons. The values are presented as percentage per dry weight. The line represents the mean and the bars SD.

**Figure 3.**
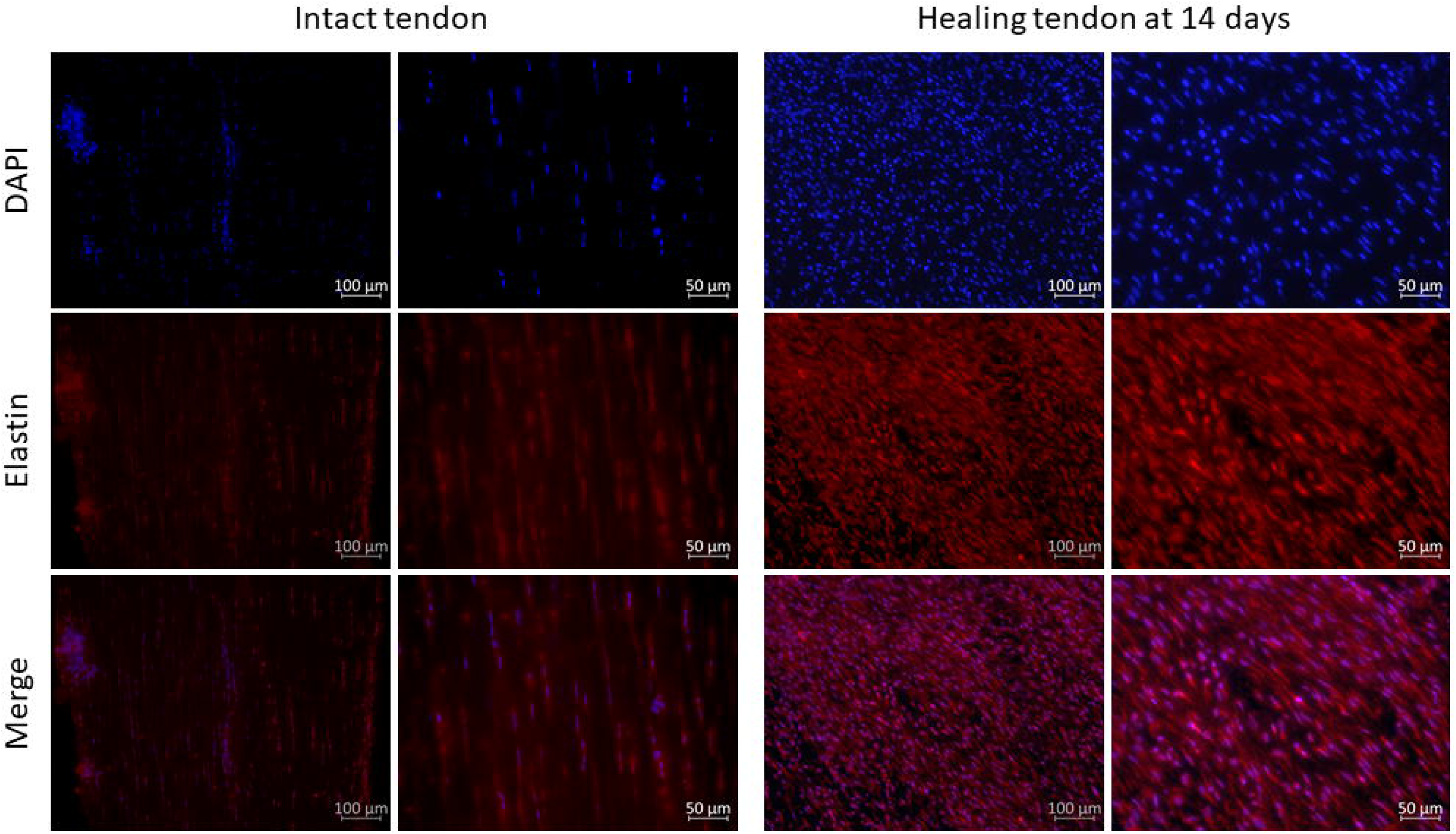
Immunohistochemical staining of elastin in intact and healing tendons. Intact and healing tendons (14 days of healing) stained with an anti-elastin antibody together with a goat anti-rabbit secondary antibody (red) and counterstained with DAPI (blue). The pictures are taken in the middle of the tendons with 20X and 40X magnification.

### Stiffness increased and creep decreased in healing tendons over time

Tendon stiffness in both the toe region and the linear region increased with time after surgery, by 90 % in the toe region (p<0.0001) and 60 % in the linear region (p<0.0001, Table 2). Albeit, there was no correlation between elastin content in the healing tendons and stiffness in either region, or stiffness ratio (simple linear regression analysis of stiffness ratio, p=0.88, R^2^=0.001, Figure 4A). Creep was significantly reduced by 30% at 28 days of healing compared to 14 days (p=0.048), while hysteresis was unaffected by time after surgery (Table 2). The transverse area was highest at day 14 and this was later reduced, 14 days later, by approximately 20% at day 28. All measured parameters were significantly different between the intact and healing tendons (p<0.001 for all).

**Table 2.** Experiment 1 - Mechanical data and elastin levels in intact and healing tendons (7, 14 or 28 days of healing).

|  | Intact tendons | Healing tendons |  |  |
| --- | --- | --- | --- | --- |
|  |  | Day 7 | Day 14 | Day 28 |
| Transverse area (mm<sup>2</sup>) | **2.8 ± 0.9** | 17.4 ± 2.4 | 21.7 ± 3.8<sup>a</sup> | 16.7 ± 2.9<sup>b</sup> |
| Stiffness toe region, 1-1.5N (N/mm) | **15.1 ± 2.2** | 3.7 ± 0.3 | 4.2 ± 0.8 | 7.0 ± 1.3<sup>a, b</sup> |
| Stiffness linear region, 7-9N (N/mm) | **30.1 ± 2.8** | 10.3 ± 0.7 | 12.5 ± 1.6<sup>a</sup> | 16.1 ± 1.7<sup>a, b</sup> |
| Stiffness ratio toe/linear | 0.5 (0.1) | 0.4 (<0.1) | 0.3 (<0.1) | 0.4 (0.1) |
| Hysteresis (%) | **8.6 ± 2.9** | 13.6 ± 1.1 | 14.3 ± 3.2 | 16.0 ± 1.6 |
| Creep (%) | **3.9 ± 0.8** | 9.7 ± 2.6 | 10.3 ± 3.7 | 7.0 ± 1.5<sup>b</sup> |
| Elastin/dry weight (%) | **2.0 ± 0.5** | 4.2 ± 1.2 | 4.1 ± 0.7 | 3.9 ± 0.6 |

Bold means significant difference ( $p \leq 0.05$ ) between intact and healing tendons tested by paired t-test. a means significantly different from day 7 and b means significantly different from day 14, tested by a one-way ANOVA. Values are expressed as mean ± SD.

**Figure 4.**
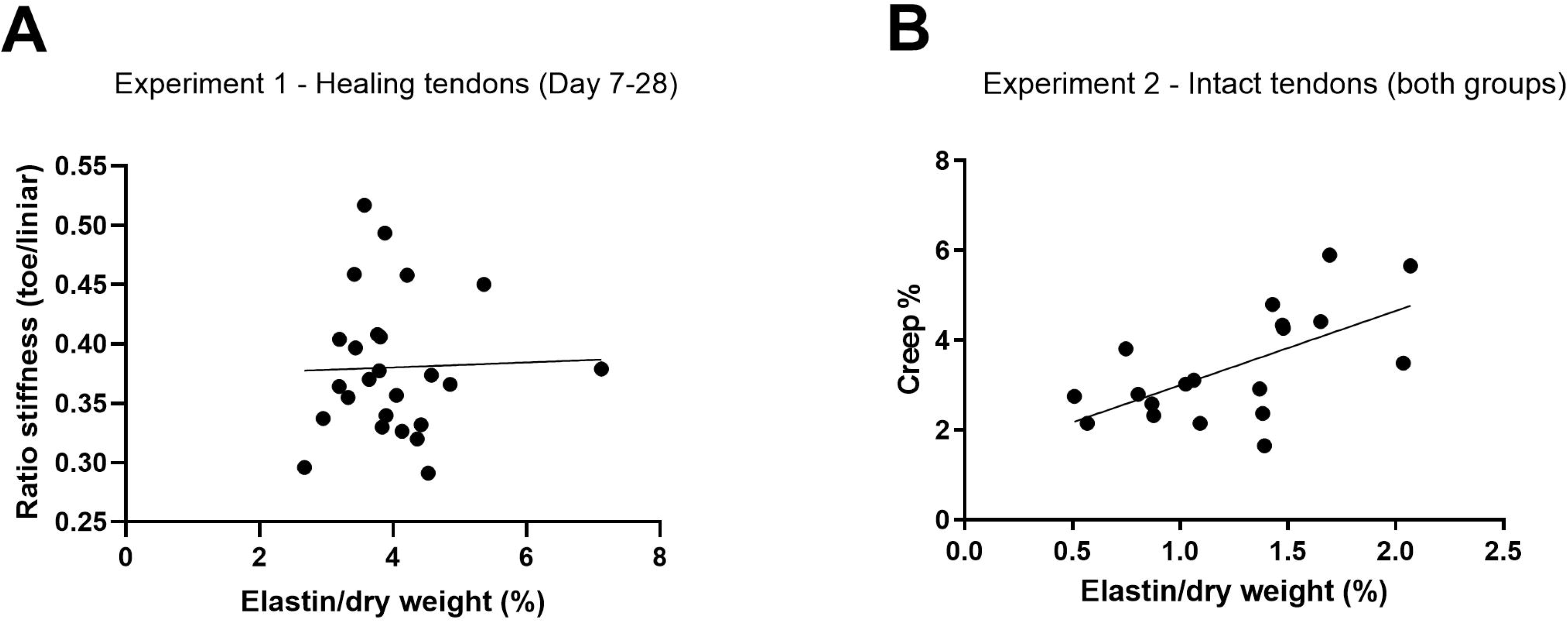
Simple linear regressions between mechanical properties and elastin levels. A) There was no correlation between stiffness ratio (toe stiffness / linear stiffness) and elastin levels of the healing tendons 7-28 days post-surgery in Experiment 1 (p=0.88, R^2^=0.001). B) There was a correlation between elastin levels and creep % of the intact tendons in Experiment 2 (p=0.005, R^2^=0.38).

### Degradation of elastin led to reduced creep in intact and healing tendons

Elastase incubation (2U/ml) for 14 hours reduced the elastin levels by 33% in intact tendons (p=0.01, Table 3) compared to controls, with a subsequent reduction by 40% in creep (p=0.001). Indeed, there was also a clear correlation between creep and elastin content in the intact tendons (simple linear regression analysis, p=0.005, R^2^=0.38, Figure 4B). Hysteresis in intact tendons, as well as stiffness in either region (toe or linear) were unaffected by elastase incubation. The creep test performed in Experiment 3 confirmed a reduction in creep, from 8.5% in the controls to 4.0% in the elastase group (p<0.001, Figure 5).

**Table 3.**
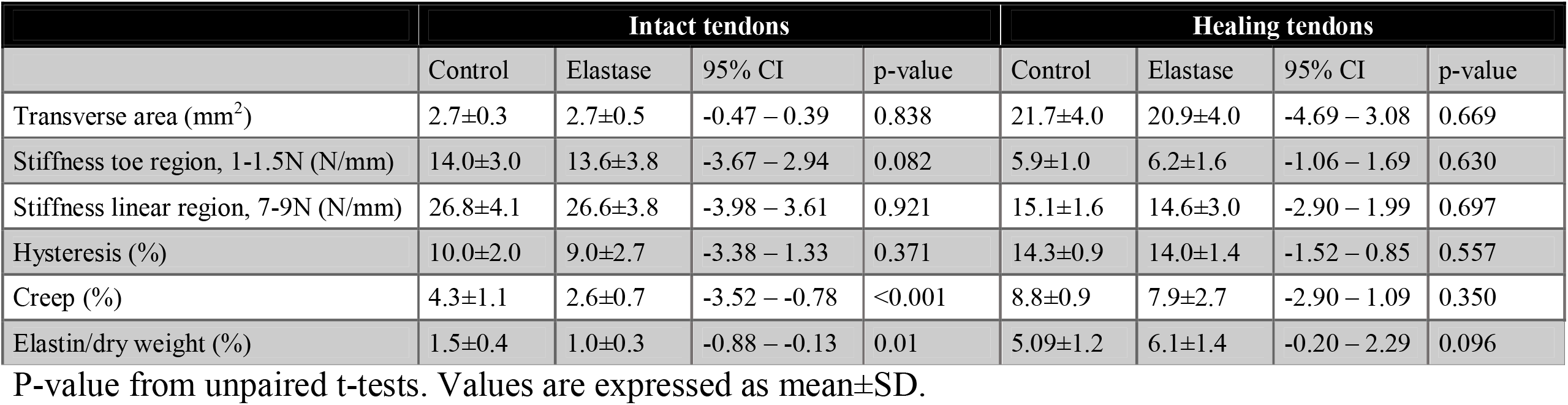
Experiment 2 - Mechanical data and elastin levels in intact and healing tendons (14 days of healing) treated with 2U/ml of elastase or PBS (controls) for 14 hours.

**Figure 5.**
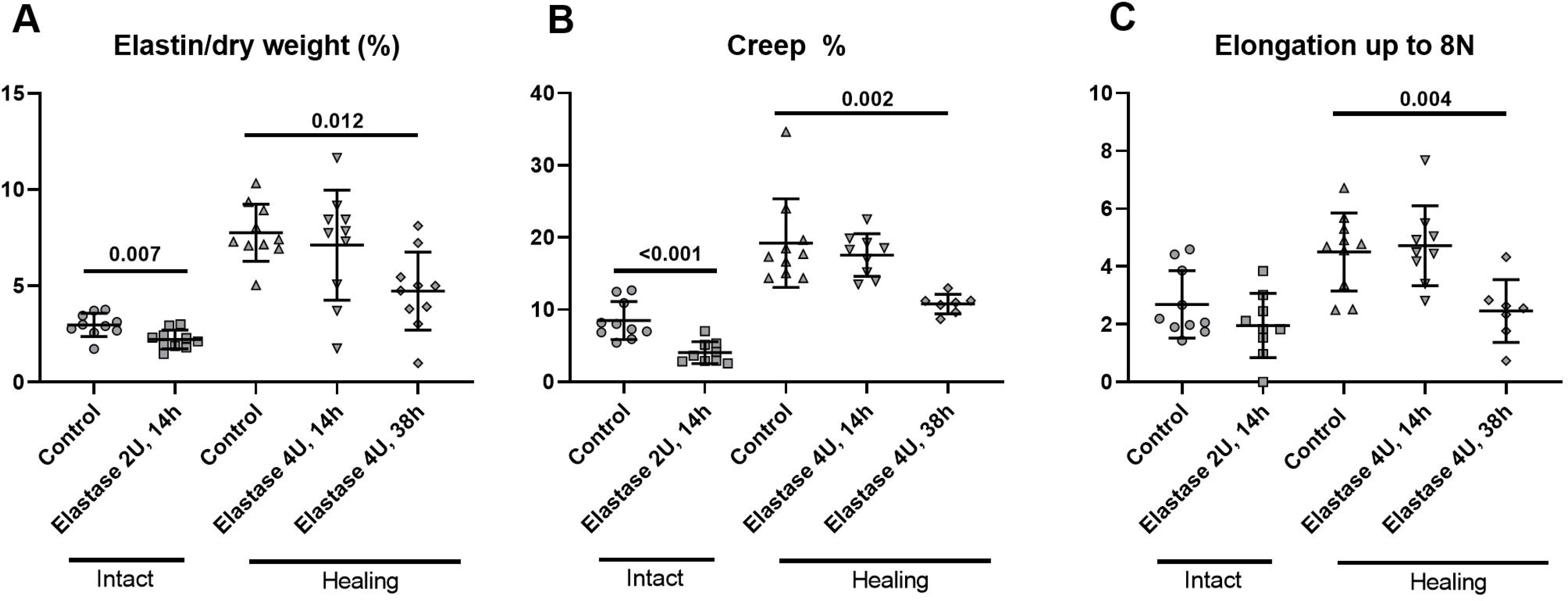
Elastin content, creep and tendon elongation up to 8N after removal of elastin. A) Elastin content (%) measured as elastin/dry weight. B) Creep (%) measured as the difference in elongation over 300 seconds. C) Elongation up to 8 N (mm) during creep test measurements. Intact and healing tendons (14 days of healing) were treated with elastase (2 or 4 U/ml) or PBS (controls) for 14 or 38 hours. Intact and healing tendons were analyzed separately with an unpaired t-test and a one-way ANOVA, respectively. All data is from Experiment 3.

Elastin levels in the healing tendons were unaffected by both 2 and 4U/ml of elastase for 14 hours and by extension there was no difference in creep between treated and untreated tendons (p=0.350, Table 4, Figure 5). However, elastase incubation with 4U/ml for 38 hours led to a 44% reduction in elastin content in the healing tendons. Creep was reduced by approximately 50% in the elastase group (p=0.002) and the tissue elongation up to the starting position for the creep test was also less in the treated healing tendons, this was not seen for the intact tendons.

**Table 4.** Experiment 3 - Mechanical data and elastin levels in intact and healing tendons (14 days of healing) treated with 2 or 4 U/ml of elastase or PBS (controls) for 14 or 38 hours.

|  | Intact tendons |  |  |  | Healing tendons |  |  |  |  |  |
| --- | --- | --- | --- | --- | --- | --- | --- | --- | --- | --- |
|  | Control | Elastase 2U/ml 14h | p-value unpaired t-test | 95% CI | Control | Elastase 4U/ml 14h | 95% CI vs control | Elastase 4U/ml 38h | 95% CI vs control | p-value ANOVA |
| Length before test (mm) | 10.1±1.8 | 12.4±2.0 | 0.012 | 0.60 – 4.14 | 13.5±1.6 | 13.6±1.8 | -1.73 – 2.00 | 16.7±2.0* | 1.31 – 5.04 | 0.001 |
| Length after test (mm) | 10.7±1.4 | 14.3±2.7 | 0.002 | 1.48 – 5.57 | 14.9±2.4 | 16.5±2.5 | -1.02 – 4.37 | 17.9±2.3* | -0.25 – 5.85 | 0.051 |
| Δ length | 0.7±1.1 | 1.5±1.5 | 0.171 | -0.41 – 2.12 | 1.4±1.2 | 3.4±1.4* | 0.63 – 3.52 | 1.0±1.2 | -1.89 – 1.11 | 0.002 |
| Gap length (mm) | - | - | - | - | 3.7±0.9 | 5.2±1.2* | 0.47 – 2.61 | 5.3±1.0* | 0.56 – 2.70 | 0.002 |
| Transverse area (mm<sup>2</sup>) | 2.7±0.6 | 3.7±1.1 | 0.02 | 0.17 – 1.81 | 25.2±4.3 | 25.7±4.5 | -5.27 – 6.30 | 23.1±7.3 | -7.89 – 3.68 | 0.543 |
| Creep start (mm) | 2.7±1.2 | 2.0±1.1 | 0.183 | -1.84 – 0.38 | 4.5±1.4 | 4.7±1.4 | -1.20 – 1.63 | 2.5±1.1* | -3.56 – -0.53 | 0.004 |
| Creep (%) | 8.5±2.6 | 4.0±1.5 | <0.001 | -6.55 – -2.33 | 19.2±6.1 | 17.6±3.0 | -6.30 – 2.97 | 10.8±1.4* | -13.4 – -3.48 | 0.002 |
| Elastin/dry weight (%) | 3.0±0.6 | 2.2±0.5 | 0.007 | -1.27 – -0.24 | 7.8±1.5 | 7.1±2.9 | -2.94 – 1.65 | 4.7±2.02* | -5.32 – 0.74 | 0.012 |

\* means significantly different from control. Values are expressed as mean±SD.

### Structural properties of the tissues were affected by elastase treatment

Effective elastase treatment (i.e. reduced elastin levels) in intact and healing tendons resulted in significantly longer tendons before the test was initiated (Table 4). Both the healing and intact tendons were also significantly longer after the creep test. The increased length was also confirmed by an increased gap length in the healing tendons treated with elastase. The transverse area was unaffected by elastase incubation in the healing tendons while there was a slight increase after elastase incubation in the intact tendons.

## DISCUSSION

The elastin levels were twice as high in healing tendons compared to intact tendons, in contrast to our hypothesis. This was confirmed by immunohistochemistry were elastin had a wider distribution throughout the extracellular matrix in the healing tendons. Elastin seems to be quickly produced during tendon healing as the levels were high already at 7 days post-surgery. Additionally, there was neither an increase nor a decrease during the following 21 days. Furthermore, the stiffness of the toe-region in relation to the stiffness of the linear region did not seem to correlate with the elastin content in the tendon. Elastin digestion by elastase had no effect on tendon stiffness or hysteresis. However, elastin appears to play a role in tendon compliance as a decreased creep was seen in tendons with reduced elastin levels.

Repair of elastic fibers has been reported to be problematic due to the complexity of the molecular assembling (17). Elastic fibers heal in an unorganized matter and this leads to improper repair and function (10). Nonetheless, our study shows high levels of elastin in healing tendons already 7 days post-surgery compared to intact tendons. This indicate that these elastin molecules are most likely newly formed elastin that possibly assembles faster than other components in tendons such as collagen. The elastin levels are thereby initially in a higher portion of the tissue dry weight, similar to healing arteries (18). Our histological analysis also showed a more general distribution of elastin in the extracellular matrix of healing tendons, in contrast to intact tendons. The intact tendons had more regular elastin distribution that were more linear and parallel to the collagen fibers. This parallel appearance resembles what others have showed in intact tendons where it has mainly been found in the interfascicular matrix and in the pericellular environment (17, 18).

Elastase treatment with 2U/ml was enough to reduce the levels of elastin in intact tendons after 14 hours of digestion. However, as the healing tendons were bigger and contained a larger amount of elastin, we needed to increase both the dose of elastase as well as the incubation time to be able to reduce the levels of elastin in healing tendons. Elastin in intact tendons has previously been described to facilitate shear stress and sliding between the fascicles (19). Another study on rat tail tendon fascicles has shown a reduction in failure strain and ultimate tensile strength (16). It is difficult to get reliable ultimate failure force data from the intact tendons as they often rupture by the clamps that is the weakest point. Hence, we did not perform any pull-to-failure test. Albeit, seven of the healing tendons in the elastase group ruptured during the cyclic and creep test. This indicates that the healing tendons might become weaker after elastase treatment similar to what was shown on the rat tail tendon fascicles (16). Moreover, the lack of change in hysteresis after elastin depletion is consistent with previously finding on medial collateral ligaments (14).

The stiffness in our healing tendons increased over time (between day 7 to 28) and this was significant for both the toe region and the linear region. We had a hypothesis that elastin content would be related to the relation between the stiffness in these two regions, as previously been described in medial collateral ligaments where the main effect was seen in the toe region (14). We could not confirm our hypothesis, and this suggest that elastin might also have an effect in the linear region for the healing tendon.

The role of elastin for viscoelastic properties in tendons has been less investigated. We showed in our study that there is a decrease in creep after elastase treatment in both intact and healing tendons. The treated tendons were also less elongated up to the initial phase of the creep test, when they were pulled up to 8 N. This suggest that elastin influence tissue compliance and this finding is consistent with results from other fibrous tissues. Degradation of elastin with elastase treatment in skin reduces its ability to deform (20). A greater tissue extensibility has also been shown in lumbar annulus fibrosus after elastase treatment (21). Elastic fibers have been suggested to be especially responsible for the recoiling mechanism after stress has been relieved (20). Perhaps a more likely scenario is that elastin affects both the recoil and the stretch mechanism. Clinical data from patients with an Achilles tendon rupture has shown that healing tendons are early on more compliable with a large amount of creep (22). This could possibly be linked to higher levels of elastic fibers and indicates that elastic fibers might be involved in a protective mechanism of the injured tendon to prevent re-injuries.

Our study showed that elastase treatment lead to an increased length in both intact and healing tendons and there was also an increased gap length. This is in line with previous studies on elastic fiber depletion where it has been shown that elastase treatment on medial collateral ligaments lead to a similar reduction in the elastin content as in our study and an increased tissue elongation (14). Furthermore, a recent study on knock-out mice for fibrillin-1, another component of the elastic fibers, described longer tendons in the deficient mice. Hence, our results on length increase after elastin disruption, together with previous data suggest that elastic fibers are involved in tendon length and possible the crimp of the collagen fibrils (23). In fact, collagen crimp has been shown to be altered after elastin disruption in rat tail tendon fascicles (16).

This study is not without limitations. Elastase has been suggested to have off target effects and it might have proteolytic activities on other compounds such as collagen, laminin, fibronectin or proteoglycans, but we did not test for that. However, this has been studied by others which could show that collagen as well as glucosaminoglycans were unaffected by elastase treatment in combination with soybean-trypsin inhibitor (15). The incubation for 14 and 38 hours probably leads to an increase in the transverse area, but the control samples were incubated in PBS, in the same condition, to overcome this (24). The elastin levels were not completely diminished but it was similar to what has previously been reported after elastase treatment on ligaments (14). However, despite this, we still had a pronounced effect on the viscoelastic properties that was measured.

In conclusion, it is well known that healing tendons are different from intact tendons regarding structural and mechanical properties and we have now showed that the differences also can be related to elastin. Healing tendons had higher levels of elastin with a more general distribution throughout the extracellular matrix compared to intact tendons. The elastic fibers appear to influence the viscoelastic properties such as tissue compliance and are possibly involved in a re-injury protective mechanism.

## ACKNOWLEDGEMENTS

The authors wish to thank Prof. Dr. Emeritus Per Aspenberg for his great contributions to design of the work and the initial analysis. Sadly, Prof. Dr. Emeritus Per Aspenberg died before the final work of the paper was conducted. This research was supported by The Swedish Research Council, The Swedish National Centre for Research in Sports, Magnus Bergvall foundation, The Swedish Society of Medicine, Linköping University and Östergötland Country Council.

## AUTHOR CONTRIBUTIONS

P. Eliasson and A. Svärd designed research; P. Eliasson and A. Svärd analyzed data; A. Svärd and M. Hammerman performed research; P. Eliasson, A. Svärd and M. Hammerman wrote the paper.

## NONSTANDARD ABBREVATIONS

PBTD: PBS containing 0.1% Tween20 and 1% DMSO

